# When should we use non-stationary adaptive management? A value of information analysis

**DOI:** 10.64898/2026.01.19.699800

**Authors:** Luz V. Pascal, Iadine Chadès, Matthew P. Adams, Kate J. Helmstedt

## Abstract

1. Making informed conservation decisions under climate change is a challenging task for practitioners, since decisions depend on changing environmental conditions and uncertain ecosystem responses to climate change. Given such uncertainties, the best practice to manage natural systems is adaptive management, where decisions dynamically adapt to the response of the ecosystem to previous conservation actions. Although adaptive approaches are optimal, they are also difficult to implement, have high computational costs, and recommend strategies that can be complex to interpret. These factors can hinder their on-ground application. On the other hand, simpler but suboptimal decision models can result in more interpretable recommendations, and might still yield good outcomes for ecosystems. Exploring trade-offs between complex optimal solutions and simpler sub-optimal solutions is essential for maximising conservation impact.
2. In this manuscript, we use value of information theory to help managers simplify their decision-models, while balancing optimality of strategies. Our approach provides modelling recommendations by determining the benefits of modelling non-stationary ecosystem dynamics and the uncertain ecosystem response to climate change. We illustrate our approach on four scenarios inspired from the management of the Great Barrier Reef, Australia, under different climate change trajectories.
3. We find that the two main drivers of the recommended reduction in model complexity are the strength of non-stationarity (e.g. climate change trajectory) and the degree of uncertainty in ecosystem responses to climate change (e.g. uncertainty in the thermal resistance of a coral reef). When non-stationarity is weak, the decision problem can be reduced from a non-stationary to a stationary formulation. Similarly, when uncertainty in the response to climate change is low, this uncertainty can be safely ignored in the decision-making process. Conversely, when non-stationarity is strong and/or uncertainty is high, our approach justifies the need to account for these complexities when making decisions, as simpler approaches would yield poor outcomes.
4. This manuscript guides managers in simplifying a modelling approach to manage ecosystems in the face of climate change. Our protocol can help simplify complex decision problems, allowing to reduce computational costs and enhance interpretability. By finding the balance between simplicity and optimality of models, this work contributes to bridging the gap between complex modelling and on-ground applications.

## 1 Introduction

Global climate change is rapidly changing environmental conditions worldwide (IPCC, 2023), exposing natural systems to unprecedented stressors. These shifts are disrupting many ecosystems’ dynamics (Williams et al., 2021), for instance by altering growth rates of fish populations (Thresher et al., 2007), reducing coral reef habitat suitability (Freeman et al., 2013) and accelerating the spread of invasive species (Keller et al., 2025). To make informed conservation decisions, practitioners must anticipate these changes and adapt their decisions accordingly (Milly et al., 2008; Nichols et al., 2011), or risk implementing inefficient management (Tucker and Runge, 2021). Research on managing non-stationary ecosystems – that is, ecosystems whose underlying dynamics are changing – is of growing interest in the literature, with applications including species translocations (McDonald-Madden et al., 2011), land acquisition (Nicol et al., 2015), natural resource management (Lindkvist et al., 2017; Tucker and Runge, 2021), and invasive species control (Keller et al., 2025).

In practice, decision-makers face two challenges when managing non-stationary ecosystems: (1) they must find strategies that are dependent on the evolving ecosystem state and changing dynamics (Tucker and Runge, 2021), and (2) they often face uncertainty about exactly how ecosystems’ underlying dynamics are changing over time, e.g. how they will respond to climate change (Dietze et al., 2018). In response to these issues, optimization approaches to manage non-stationary and uncertain ecosystems use adaptive management and are already established in the ecology literature (McDonald-Madden et al., 2011; Nicol et al., 2015; Lindkvist et al., 2017). Adaptive management enables the management of uncertain ecological systems (Walters and Hilborn, 1978; Keith et al., 2011) by dynamically adapting recommendations to the perceived effects of decisions. In non-stationary contexts, adaptive management has provided recommendations for species translocation under climate change (McDonald-Madden et al., 2011), the management of migratory shore-birds under uncertain sea level rise (Nicol et al., 2015) and the harvest of fisheries in warming oceans (Lindkvist et al., 2017). These recommendations depend upon the dynamically changing ecosystem state as well as extrinsic factors affecting those dynamics like progressive climate change.

Despite these examples, in most cases applying adaptive management approaches in non-stationary environments remains challenging. Finding optimal adaptive management strategies generally requires solving complex sequential decision problems, often framed as a Partially Observable Markov Decision Process (POMDP) (Chadès et al., 2017), an optimization model for stochastic sequential decision-making (Åström, 1965; Chadès et al., 2021). POMDPs are complex decision problems to solve (Papadimitriou and Tsitsiklis, 1987; Madani et al., 1999), typically requiring approximate solutions (Kurniawati et al., 2008). A major challenge of using POMDPs in ecology is the difficulty of interpreting solutions and providing clear recommendations to managers (Dujardin et al., 2015, 2017; Pascal et al., 2020; Ferrer-Mestres et al., 2021), which limits their uptake. Explicitly including a time variable as needed in non-stationary settings further increases computational costs and reduces interpretability.

It is therefore essential to identify when fully optimized non-stationary adaptive management approaches are methodologically justified. In some applications, simpler suboptimal approaches may still yield good outcomes for ecosystems at lower computational and interpretability costs. Here, we use the value of information theory (Raiffa and Schaifer, 1961; Canessa et al., 2015) to guide decision-makers in simplifying their decision-models for managing non-stationary ecosystems under uncertainty. The value of information is often used to quantify the expected gains in benefits when uncertainties are resolved (Raiffa and Schaifer, 1961; Canessa et al., 2015). Here, we use it quantify the benefits of accounting for the uncertain ecosystem response to climate change and the non-stationarity of their dynamics. Applications of the value of information in ecology include species translocation when success is uncertain (Canessa et al., 2015; Akinlotan et al., 2024), finding trade-offs between managing and monitoring species (Maxwell et al., 2015), and evaluating the benefits of understanding feedbacks within ecosystem motifs (Xiao et al., 2019). While value of information can indicate whether an adaptive management approach is suitable (Box 1, Fig. 1 of Chadès et al., 2017), it remains unclear how to adapt this approach to non-stationary contexts.

**Figure 1:**
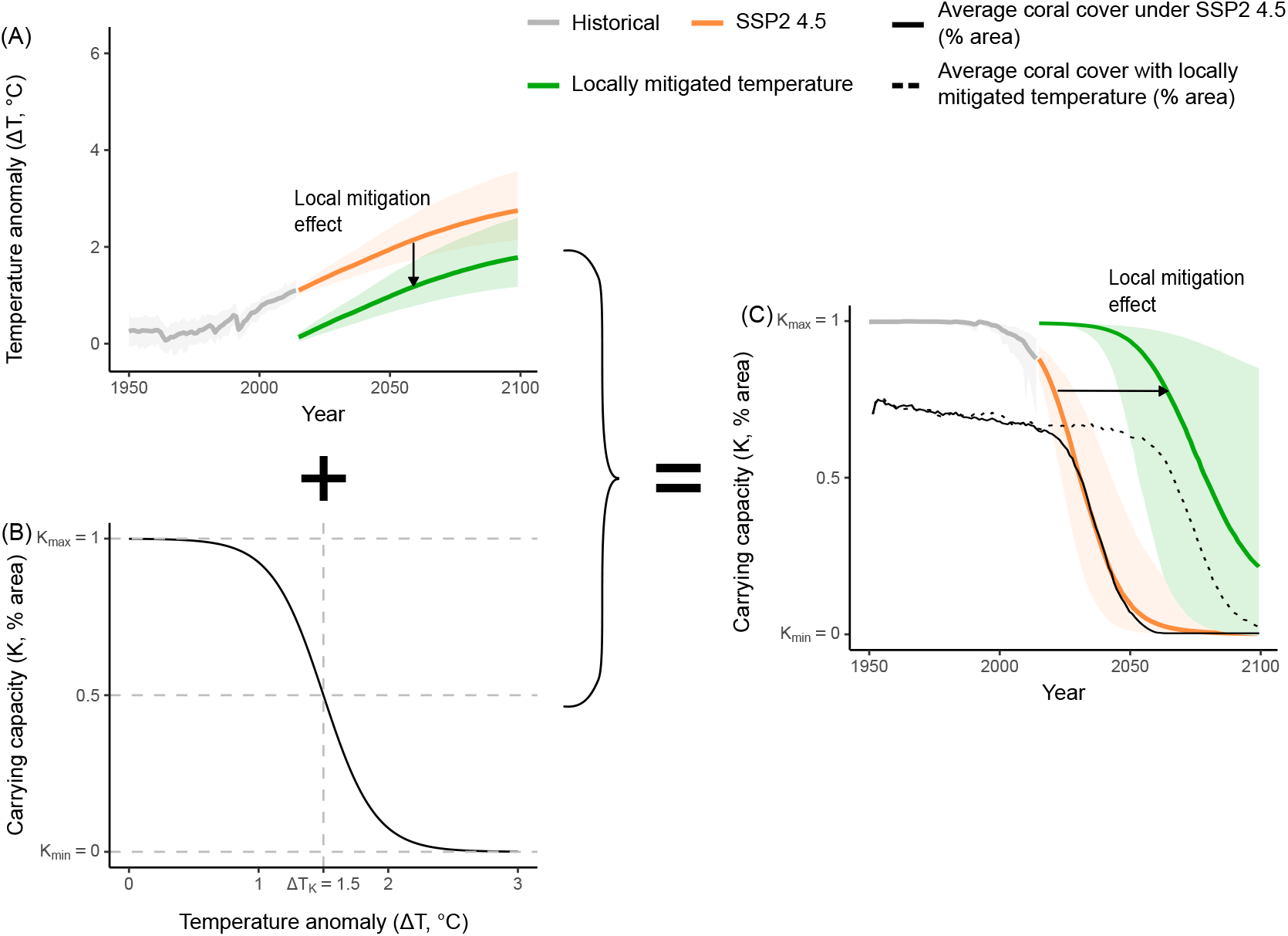
**A.** Historical and projected global mean surface air temperature anomalies relative to 1850-1900 (ΔT), according the climate trajectory SSP2 4.5. The green line represents the local mean surface air temperature anomalies when the manager locally mitigates the temperature (here by ΔT^tech^ = 1° C). **B**. Modelled effect of temperature anomalies (ΔT) on carrying capacity (*K*). Here the temperature anomaly tipping point ΔT_*K*_ is 1.5°C. **C**. Resulting modeled shifts of the carrying capacity for the climate trajectory SSP2 4.5, with and without local temperature mitigation. The solid and dotted black lines represent the average coral cover with and without local temperature mitigation (in % area).

Here, we build on these approaches to develop a value of information protocol for simplifying non-stationary adaptive management problems. Our approach finds the best trade-off between simplicity and optimality of optimization approaches. We design a new metric to quantify the benefits of modelling non-stationary dynamics and adapt the work of Xiao et al. (2019) to assess the value of knowing the ecosystem response to climate change.

We demonstrate our approach on the management of a coral reef ecosystem under changing environmental conditions (i.e., non-stationary). Worldwide, coral reefs are exposed to growing pressures including ocean acidification (Hoegh-Guldberg et al., 2017), increased frequency and severity of heat waves (Smale et al., 2019), and outbreaks of coral predators such as the Crown-of-thorns starfish (Rotjan and Lewis, 2008). In Australia, innovative solutions are currently being designed to help reefs of the iconic Great Barrier Reef cope against these stressors, with examples including genetically enhancing corals, applying cooling and shading techniques and implementing the bio-control of invasive species and coral predators (Mead et al., 2019). Planning the development and deployment of these new technologies is essential to make the most informed decisions, however it is unclear whether the non-stationarity of the ecosystems should be accounted for (Anthony et al., 2020; Pascal et al., 2025).

## 2 Methods

### 2.1 Methods overview

We introduce our value of information approach through an illustrative example inspired from the management of the Great Barrier Reef using new technologies. We consider a coral reef threatened by increasing global temperatures, which are changing the dynamics of this ecosystem over time. A manager of this coral reef has access to a new technology capable of locally mitigating the temperature to preserve the reef. The manager must decide at each discrete time-step (every year) the intensity of local temperature mitigation, balancing costs and benefits. However, the manager is uncertain about the ecosystem response to climate change. The decisions of the manager must thus depend on this uncertainty and on the changing dynamics of the system, which can be achieved using non-stationary adaptive management. Motivated by this decision-problem, we design a value of information approach, which enables the manager to find the trade-off between the optimality of non-stationary adaptive management and the simplicity of other approaches.

For ease of presentation, we first model this problem as a non-stationary Markov Decision Process (MDP, Sigaud and Buffet 2013; Marescot et al. 2013), assuming first that the decision-maker is certain about the ecosystem response to climate change. This non-stationary MDP enables the decision-maker to adapt their decisions to projected changes in temperature. Then, we detail a non-stationary adaptive management approach, allowing the manager to simultaneously account for the future projected temperatures and the uncertain ecosystem response to climate change. Finally, we present our value of information approach and demonstrate how to apply it.

### 2.2 Non-stationary Markov Decision Process model for managing a population under climate change: known ecosystem response to climate change

#### 2.2.a Ecosystem states and dynamics

We model the dynamics of coral cover (i.e., the percentage of the sea floor covered by coral) of a coral reef ecosystem. The coral cover, described by the state variable *s*_*t*_, changes at discrete time steps (*t*) according to a logistic growth model (Murray, 2002) (Equation (1)). At low abundance, the coral cover growth is approximately exponential (representing low competition for resources), and at higher abundances the coral cover plateaus as it approaches the carrying capacity (representing high competition for resources). The change in coral cover has a stochastic component, which we model using a log-normal distribution following McDonald-Madden et al. (2011). The ecosystem dynamics are then:

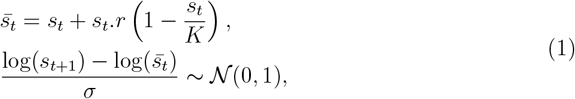

where *r* is the intrinsic growth rate (year^−1^), *K* is the carrying capacity (% area) and *σ* is the standard deviation. We initially set *r* = 0.2, *σ* = 0.2 and we assume that the the carrying capacity is negatively impacted by climate change, as described in Section 2.2.b.

#### 2.2.b Climate change effects on ecosystem dynamics

Climate change is inducing an increase of global mean surface air temperature, which is degrading coral reefs. These temperature anomalies are typically reported as relative to 1850-1900 (pre-industrial time period, IPCC 2023) and we denote ΔT_*t*_ as the temperature anomaly at year *t*. The severity of temperature changes will depend on collective efforts to mitigate climate change (IPCC, 2023), and various climate change trajectories are explored by climate scientists (Figures 1 and S1).

We incorporate the negative impact of climate change on the coral reef population into our model through changes in the coral cover carrying capacity. We assume that increases in sea surface temperature reduces carrying capacity in a non-linear relationship, such that, as temperatures increase and the carrying capacity declines, coral cover decreases, which might lead to collapse (Figure 1C).

We model a carrying capacity *K* that is logistically dependent on the temperature anomalies (ΔT, Figure 1B):

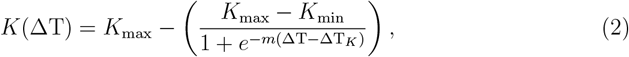

where *K*_max_ is the carrying capacity (in % area) when the temperature anomaly is low, *K*_min_ is the carrying capacity (in % area) when the temperature anomaly is high, *m* is a shape parameter (°C^−1^) controlling how the carrying capacity shifts with the temperature. We set *K*_max_ = 1, *K*_min_ = 0 and *m* = 5.

The inflection temperature ΔT_*K*_ represents the ecosystem response to climate change and is an indicator of the reef thermal resistance. This is because the higher ΔT_*K*_, the higher the temperature at which the carrying capacity declines, and the later in time the ecosystem collapses in response to gradual climate change. In a non-stationary MDP model, ΔT_*K*_ is known by the decision-maker, but later in this study, we will assume it is uncertain and can range between 1° C and 2.5° C.

#### 2.2.c Set of actions: new technologies to prevent climate change impacts

To preserve the reef ecosystem, we assume that a local manager of the reef has access to a new technology that locally mitigates the effects of climate change and aims to preserve the reef ecosystem (Figure 1A), such as water cooling and shading technologies described in Mead et al. (2019). At each time step, the manager must decide on the intensity of temperature mitigation: by 0, 1 or 2°C, denoted ΔT^tech^. This local temperature mitigation can help delay the decline of the carrying capacity, and thus delay the collapse of the ecosystem (Figure 1C). The carrying capacity then follows:

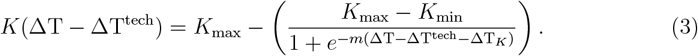

#### 2.2.d Reward function

The ecosystem provides some benefit at each timestep, which depends upon the ecosystem state (i.e., coral cover). Similarly, the manager must pay a cost to manage the ecosystem which depends on the action chosen. We combine these benefits and costs to design a reward function with two components. The manager aims to maximise the discounted sum of these rewards over the entire management time horizon. The reward achieved is proportional to the coral cover *s*, and deployment costs depend on the chosen temperature mitigation intensity. Here we assume cost is proportional to the square of the temperature mitigation intensity (ΔT^tech^), so that large interventions are linearly more expensive than small ones (Lapeyrolerie et al., 2022). The reward function is then:

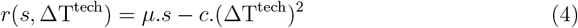

where we assume that the proportionality constant required in this reward function take values *µ* = 1 and *c* = 0.05.

#### 2.2.e Objective function

The overall objective of the manager is to find a strategy (*π*) that maximizes long term benefits. Here, the manager aims to maximize the overall ecological benefits over a finite time period (from 2015 until 2100, corresponding to the IPCC temperature projections), placing greater value on recent times compare to future ones. This can be achieved by maximizing the expected sum of discounted rewards over a finite horizon:

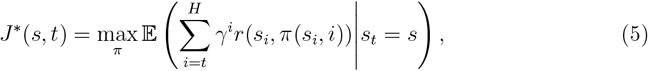

where *H* is the horizon (year 2100) and *γ* is the discount factor, relating the importance of immediate rewards compared to future ones.

In this MDP example (when we assume that the ecosystem response to climate change is fully known i.e. known value for ΔT_*K*_), an optimal strategy is a mapping from the set of states and time steps to the set of actions (Sigaud and Buffet, 2013; Marescot et al., 2013). We solve this non-stationary MDP over an 85-year time horizon (from 2015 until 2100) using the value iteration function of the MDPtoolbox (Chadès et al., 2014), and we set the discount factor *γ* = 0.99 (Section A).

### 2.3 Non-stationary adaptive management as a POMDP

In practice, the future thermal resistance of the reef is unknown to decision-makers (uncertain and unobservable value of ΔT_*K*_). As a result, MDP solutions which assume a known thermal resistance value may provide poor recommendations.

To account for the uncertain ecosystem response to climate change, we can frame the adaptive management problem by augmenting the MDP (Section 2.2) with a set of non-observable states. These non-observable states represent alternative possible ecosystem responses to climate change (represented by alternative inflection temperature values ΔT_*K*_), with one assumed to be correct but unknown to the decision-maker. To capture this uncertainty, the decision-maker has a belief (a probability distribution) over the non-observable states (denoted *b*), representing the probability that each alternative inflection temperature is correct. The belief updates following Bayes’ rule after each action and observed coral cover value, which allows the decision-maker to sequentially adapt their decisions (see Appendix for details). Following these steps converts the problem into a Partially Observable MDP (POMDP, Chadès et al., 2012, Section B)

### 2.4 Value of information protocol

#### 2.4.a Protocol overview

A non-stationary adaptive management approach explicitly accounts for non-stationary and uncertain ecosystem dynamics. Specifically, non-stationarity requires accounting for the time dimension and uncertainty in the ecosystem dynamics requires using probabilistic a state-action transition function instead of deterministic state-action transition. While using non-stationary adaptive management is guaranteed to be optimal and to generate the highest achievable benefits, it also comes at high computational costs and produces recommendations that are challenging to interpret (Dujardin et al., 2015, 2017; Ferrer-Mestres et al., 2021). Especially in Ferrer-Mestres et al. (2024), authors demonstrate that increasing dimensions makes it challenging to interpret the solutions of MDPs. Removing one or both of these complexities simplifies the model, enhances interpretability and reduces computational costs, resulting in an approach that is more likely to be applied in practice. However, this simplification also results in a loss of optimality and a reduction in obtained benefits (i.e. loss in the cumulative total benefits Section 2.2.e).

The objective of our protocol is to rapidly estimate this loss of benefits, before solving a non-stationary adaptive management problem. As detailed in Sections 2.4.c and 2.4.d, this calculation also identifies the best simplified strategy that can be achieved by reducing the complexity of the decision model. In this way, our protocol helps decision-makers weigh whether they are willing to lose some benefits in exchange for simpler and more interpretable model and strategy. To determine this trade-off, decision-makers must choose an acceptable threshold of expected losses (*δ*). This threshold represents the maximum proportion of total benefits they are willing to trade for a simpler decision model (e.g., up to 1% loss of total benefits compared to the fully complex model in exchange of a simple stationary MDP).

After defining the threshold, we calculate these expected losses using the value of information theory, which evaluates whether a gain of information improves expected benefits (Section 2.4.b). For each complexity (uncertainty and non-stationarity), we provide a value of information metric (EV_model_ and EV_non−stat_, see Sections 2.4.c and 2.4.d). These metrics quantify the expected loss in performance due to the simplification of the model by removing that complexity. Each metric thus answers one of the following questions:

- ***Q1***. *Does accounting for the uncertainty in the ecosystem response to climate change improve decision-making outcomes?*
- ***Q2***. *Does accounting for non-stationarity of the ecosystem dynamics improve decision-making outcomes?*

If the value of a metric is lower than the predefined threshold of acceptable losses (*δ*), the decision-model can be simplified – in the best case from a non-stationary adaptive management problem as a POMDP to a simple stationary MDP. The possible outcomes of the protocol are summarized in Table 1.

**Table 1:** After calculating the value of information metrics, and compared with the acceptable loss threshold, we have four outcomes (corresponding to the four possible combinations of answers to questions Q1 and Q2). We summarize the possible outcomes of this analysis in this table. The value of accounting for model uncertainty EV_model_ is defined in Section 2.4.c and the value of modelling non-stationarity is defined in Section 2.4.d.

| Value of Information protocol |  | **Q1. Does accounting for the uncertainty in the ecosystem response to climate change improve decision-making outcomes?**<br>**i.e. is $EV_{\text{model}} \geq \delta$?** |  |
| --- | --- | --- | --- |
|  |  | No | Yes |
| **Q2.**<br>**Does accounting for non-stationarity improve decision-making outcomes ?**<br>**i.e. is $EV_{\text{non-stat}} \geq \delta$?** | No | Stationary Markov Decision Process (MDP)<br>(e.g. Marescot et al. (2013)) | Stationary adaptive management<br>(e.g. Chadès et al. (2012), Fackler and Pacifici (2014), Pascal et al. (2025)) |
|  | Yes | Non-stationary MDP<br>(e.g. Tucker and Runge (2021), Keller et al. (2025)) | Non-stationary adaptive management<br>(e.g. McDonald-Madden et al. (2011), Nicol et al. (2015), Lindkvist et al. (2017)) |

We now detail how to apply our protocol in practice, by introducing the value of information theory and the two metrics needed to answer **Q1**. and **Q2**.. We demonstrate our approach on four case studies (Section 2.5), corresponding to the four possible combinations of answers to these questions.

#### 2.4.b The relative Expected Value of Perfect Information

The two metrics used in our protocol are value of information metrics (Raiffa and Schaifer, 1961; Canessa et al., 2015; Xiao et al., 2019). Traditionally, the value of information quantifies the expected improvement in outcomes from additional information, which here we equivalently use to calculate the expected loss of benefits due to the absence of information. We use this framework to estimate the importance of accounting for the uncertainty in the ecosystem response to climate change and the non-stationarity of its dynamics.

Importantly, our value of information approach allows us to estimate potential losses in benefits without solving the full non-stationary adaptive management problem. We do so by adopting the *no-learning* interpretation of the value of information (Chadès et al., 2017). Under a *no-learning* setting, the value of information is computed assuming that, in the presence of uncertainty, the management strategy is fixed in advance. Consequently, the action taken (temperature mitigation) in each state (i.e. coral cover - year pair) is determined at the outset of the decision problem and does not change in response to new information.

As we show in Sections 2.4.c and 2.4.d, under in a *no-learning* setting, we only need to solve MDPs, which makes this interpretation computationally more tractable than the alternative interpretation proposed by Williams and Johnson (2015). In the latter interpretation, the value of information is calculated by allowing the strategy under uncertainty to update as new information is obtained, which for our problem corresponds to solving the full non-stationary adaptive management problem (Williams and Johnson, 2015; Chadès et al., 2017).

Our two metrics measure the relative Expected Value of Perfect Information (rEVPI; Xiao et al. 2019), that is the relative difference between the expected outcomes of an optimal strategy under perfect information (EV_certainty_) and the expected outcomes of an optimal strategy under uncertainty (EV_uncertainty_):

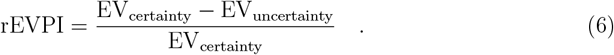

As a clarification, here the value of information measures the improvement in expected management outcomes. As strategies with different decisions, might still result in the same outcomes, changes in decisions are not good indicators of the value of information (Akinlotan et al., 2024).

We now introduce the two rEVPI-based metrics, EV_model_ and EV_non−stat_.

#### 2.4.c Q1. Does accounting for the uncertainty in the ecosystem response to climate change improve decision-making outcomes? The value of accounting for model uncertainty

To answer **Q1**, we apply the value of information approach of Chadès et al. (2017) and Xiao et al. (2019) to quantify the expected loss of benefits from removing the uncertainty in the ecosystem response to climate change (ignoring uncertainty about the true value of ΔT_*K*_). We refer to this metric as the *value of accounting for model uncertainty*. The manager perfectly knows the climate change trajectory (current SSP), but they are uncertain about ΔT_*K*_. Instead, they have a discrete set of possible scenarios (alternative possible values for ΔT_*K*_), that we denote *Y*, and a prior *b*_0_ over these scenarios, where *b*_0_(*y*) is the probability that the scenario *y* is correct.

The value of accounting for model uncertainty is:

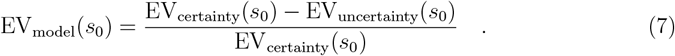

EV_certainty_ is the expected outcome that could be achieved if the uncertainty about ΔT_*K*_ were resolved. It is the weighted average of the maximum expected benefits for each scenario in *Y* :

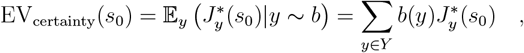

where 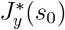is the maximum achievable expected sum of discounted rewards when the true scenario is *y* and the initial state is *s*_0_ (the value function of MDP *y*). Calculating EV_certainty_(*s*) requires solving |*Y* | non-stationary MDPs (Section A).

Under uncertainty, the decision maker must choose a strategy without knowing the ecosystem’s response to climate change (i.e. the value of ΔT_*K*_), aiming to maximize benefits across all possible models in *Y*. We apply the method of Xiao et al. (2019) to compute EV_uncertainty_, by identifying the best strategy, when no learning occurs (i.e. a strategy of the form *π* : *S* × Θ → *A*). As highlighted by the authors, finding this optimal strategy is a difficult combinatorial problem (with |*A*|^|*S*||*H*|^ = 3^11×85^ possible combinations). Instead, Xiao et al. (2019) estimate a lower bound of EV_uncertainty_, by evaluating a smaller set of strategies, denoted Π_Y_. This set Π_Y_ consists of the optimal strategies for each individual model in *Y*. EV_uncertainty_ is then approximated as:

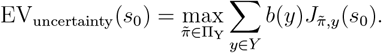

Calculating EV_uncertainty_ involves first solving each MDP in *Y* and storing its optimal strategy in Π_Y_. For every strategy 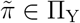, we then compute its value 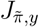 when applied to each model *y* ∈ *Y*. These values are averaged using the prior distribution *b*. EV_uncertainty_ is the highest expected value among all strategies in Π_Y_.

The best no-learning strategy is then:

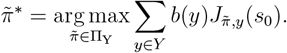

Importantly, if EV_model_ falls below the decision-maker’s predefined threshold (*δ*), the simpler strategy 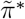 achieves nearly optimal outcomes. Otherwise, our protocol recommends implementing full adaptive management.

#### 2.4.d Q2. Is non-stationarity valuable for improving decision-making outcomes? The value of modelling non-stationarity

To answer **Q2**., we build on Tucker and Runge (2021) and Xiao et al. (2019) to propose a new value of information measure: the value of non-stationarity, that quantifies the benefits of modelling non-stationary dynamics. We define this metric as the expected loss in benefits resulting from applying a stationary strategy to a non-stationary system. For any non-stationary MDP *y*, we define the value of modelling non-stationarity as:

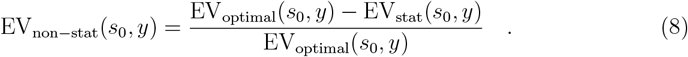

EV_optimal_(*s*_0_, *y*) represents the maximum achievable expected benefits when the decision-maker accounts for the non-stationary dynamics of the system, and the initial ecosystem state is *s*_0_. It corresponds to the optimal value function of the non-stationary MDP *y*. Therefore:

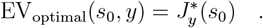

EV_stat_(*s*_0_, *y*) represents the maximum expected benefits achievable when applying a stationary strategy to the non-stationary system *y*, given the initial state *s*_0_. Assuming the manager selects the stationary strategy that yields the best outcomes, we define:

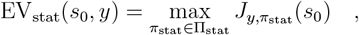

where Π_stat_ is the set of all possible stationary strategies, defined as mapping only from the set of states to the set of actions (independent of time, *π*_stat_ : *S* → *A*). For each *π*_stat_ ∈ Π_stat_, 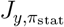 (*s*_0_) represents the expected benefits of applying *π*_stat_ to the non-stationary MDP *y*.

Similarly to the value of model uncertainty (Section 2.4.c), finding the best stationary strategy is a combinatorial problem, with |*A*|^|*S*|^ = 3^11^ possible combinations (for each state *s* there are |*A*| possible actions). Instead of testing all possible strategies, we estimate a lower bound of EV_stat_ by building on Xiao et al. (2019) and selecting a subset of stationary strategies that are likely to yield high benefits. Specifically, we evaluate three stationary strategies: *π*_init_, *π*_avg_ and *π*_final_. We obtain each strategy by solving a stationary MDP under constant temperature anomalies: (1) the initial temperature in 2015 (*π*_init_), (2) the average temperature between 2015 and 2100 (*π*_avg_) and (3) the final temperature in 2100 (*π*_final_). We then apply each strategy to the non-stationary MDP *y* and select the one that yields the highest benefits. Thus, for our example, Π_stat_ = {*π*_init_, *π*_avg_, *π*_final_}.

Similarly to EV_model_, calculating EV_non−stat_ helps identify the best-performing stationary strategy:

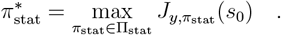

Decision-makers may choose to implement this stationary strategy or not depending on the value of EV_non−stat_, giving the expected loss in benefits compared to the fully informed optimal strategy.

### 2.5 Details of the case studies

We design four case studies summarized in Table 2 to illustrate our approach, and we select these case studies intentionally to match the four possible outcomes of our protocol (Table 1). For each case study, we select two possible ecosystem responses to climate change (two values for ΔT_*K*_) corresponding to low and high thermal resistance scenarios of the ecosystem. In the low thermal resistance scenario (lower ΔT_*K*_), ecosystem decline begins at lower temperatures, while in the high resistance scenario, decline occurs at higher temperatures. The decision-maker is uncertain which scenarios is correct. We set an acceptable loss threshold of *δ* =1% and apply our protocol to guide the selection of the modelling approach. We then extend our analysis by generalizing the results across all ΔT_*K*_ between 1°C and 2.5°C.

**Table 2:**
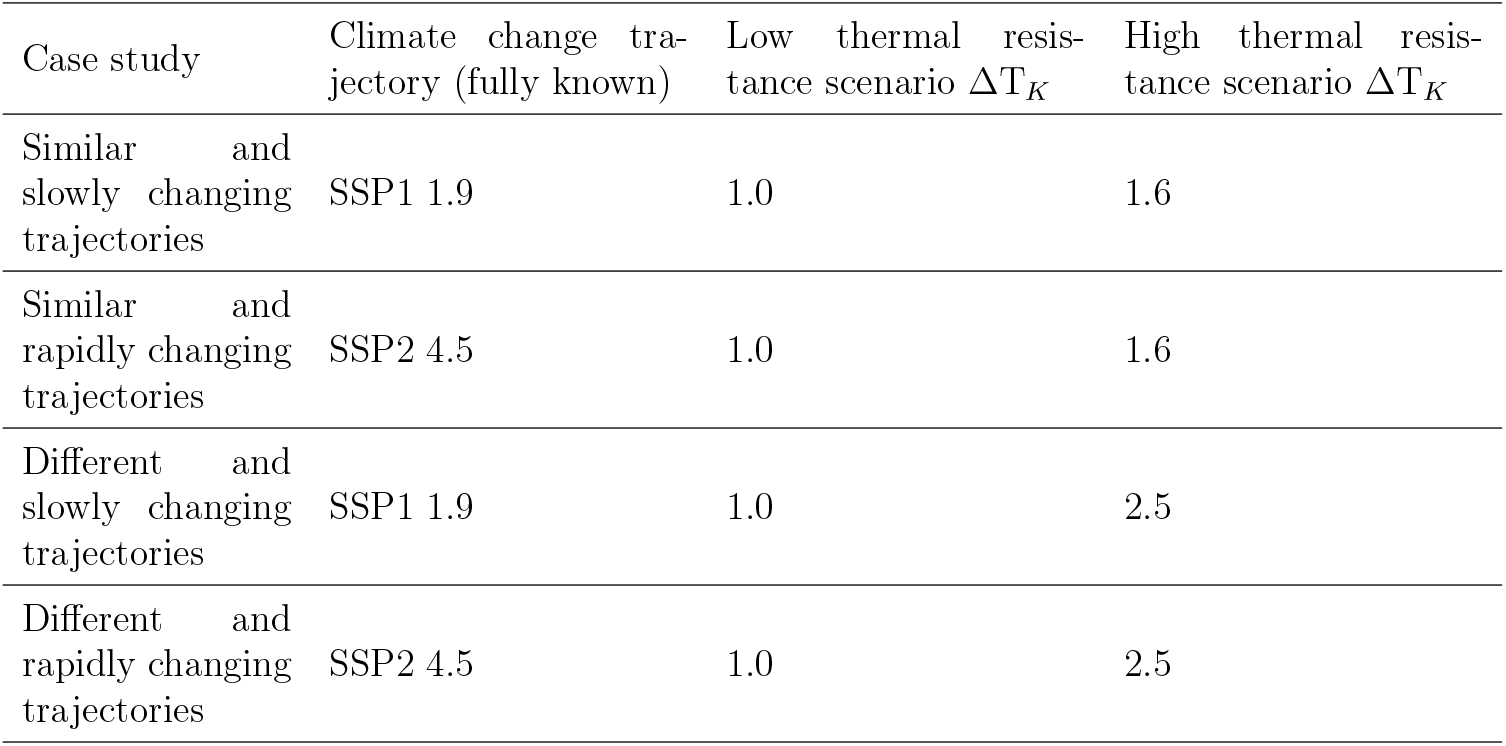
Summary of four case studies. We use the temperature anomaly projections of IPCC (2023) for each climate trajectory (Figure S1).

## 3 Results

### 3.1 Four case studies

Our value of information approach allows to quantify the trade-off in strategy performance between the optimality of a non-stationary adaptive management program and the simplicity of a stationary MDP. We now detail on the four selected case studies that we intentionally chose to match the four possible outcomes of our protocol (Table 1).

In a first case study, that has similar and slowly changing trajectories of the carrying capacity (Figure 2A), the carrying capacity in both the low and high thermal resistance scenarios decline without technological intervention (ΔT^tech^ = 0). These two trajectories for the carrying capacity differ by around 20%, but their decline is mild (less than 10% difference between the start and end values, Figure 2A(i)). This similarity between the trajectories results in a low value of resolving model uncertainty (0.1%) and the slow decline yields a low value of modelling non-stationarity (0.1% and 0.2% for the low and high thermal resistance respectively). These value of information metrics are below the acceptable loss threshold of *δ* =1%, indicating that the decision-maker can simplify the decision problem from a non-stationary adaptive management problem to a stationary MDP, with the guarantee of losing less than 1% of total benefits. The recommended simpler strategy is to locally mitigate temperature by 1°C throughout the entire decision period, provided coral cover remains positive. In this case study, the computational benefit of using a stationary MDP drops the computational time from 2h to solve a non-stationary adaptive management problem to 0.2 s.

**Figure 2:**
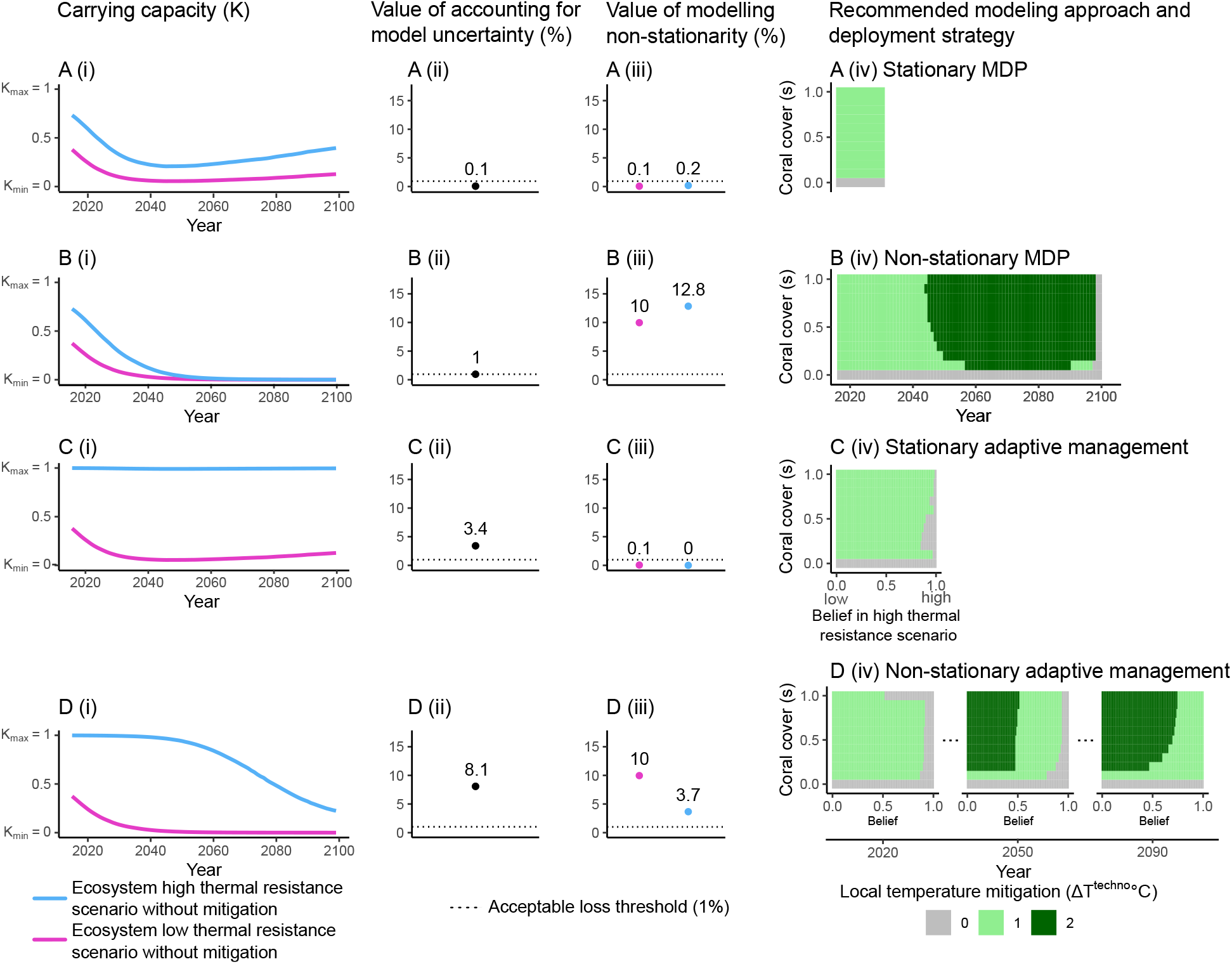
Illustration of how our protocol is applied in practice, using the four case studies of Table 2. For all case studies, the first column represents two carrying capacity trajectories: one for high thermal resistance (higher ΔT_*K*_) and one for low thermal resistance (lower ΔT_*K*_) without temperature mitigation. The second column represents the value of accounting for model uncertainty. The third column depicts the value of modelling non-stationarity for each trajectory of the carrying capacity. The fourth column shows the recommended model simplification and deployment strategy (outcome of our protocol).

In a second case study, where the carrying capacity trajectories are similar and rapidly changing (Figure 2B), the two alternative carrying capacity trajectories follow a similar decline but, unlike case A, drop to zero around 2050. This structural change significantly increases value of non-stationarity (increased between 10% and 12.8%), while the value of resolving model uncertainty remains low (1%). Our protocol recommends using a non-stationary MDP. Solving this non-stationary MDP only requires 2s of computational time, compared to 2h for non-stationary adaptive management. The recommended strategy is to locally mitigate the temperature by 1°C for almost all ecological states from year 2015 until 2050 approximately, and then intensify mitigation to 2°C until the end of the decision period (Figure 2B(iv)). The deployment stops in 2100, due to the finite time horizon.

For the third case study, with different and slowly changing trajectories (Figure 2C), the two trajectories of the carrying capacity are very different, but seem almost constant through time. Our analysis demonstrates that the most valuable variable for decision-making is the uncertainty in the ecosystem response to climate change: the value resolving this uncertainty is 3.4% (Figure 2C(ii)), while the value of non-stationarity is negligible (0-0.1%, Figure 2C(iii)). These results suggest that a stationary adaptive management approach can still yield high benefits for the ecosystem. The recommended strategy is to mitigate temperature by 1°C for most beliefs, but to stop mitigation under the highest beliefs in the high thermal resistance scenario.

For the fourth case study, with different and rapidly changing trajectories (Figure 2D), the carrying capacity trajectories differ greatly and are time-dependent, with early decline in the low thermal resistance scenario and late decline in the high resistance one. Both the uncertainty in the ecosystem response and the non-stationarity significantly affect decision-making, with a value of accounting for model uncertainty reaching 8.1%, and the value of modelling non-stationarity between 3.7% and 10%. This shows that removing either complexity would lead to substantial losses, making a simpler optimization models inadequate. The recommended model is a non-stationary adaptive management program. The optimal strategy depends on the coral cover, the time step, and the belief in the thermal resistance scenarios. Around 2020, it is optimal to mitigate temperature by 1°C for most belief states, except if the belief in the high thermal resistance scenario exceeds 0.9, where no mitigation is needed. By 2050, if the belief is below 0.5, intense mitigation of 2°C is recommended; for beliefs between 0.5 and 0.9, moderate mitigation of 1°C remains optimal; above 0.9, no mitigation is recommended. Toward the end of the decision period, intense mitigation (2°C) is optimal if the belief is below 0.6, and moderate mitigation (1°C) if above.

### 3.2 The value of accounting for model uncertainty

When the decision-maker is uncertain between two ecosystem responses to climate change (i.e. two possible values for ΔT_*K*_), the value of model uncertainty depends on both the difference between these values and the climate change trajectory (Figure 3). Generally, the greater the difference between the two ΔT_*K*_ values, the higher the model uncertainty. This effect is the most pronounced under the most pessimistic climate trajectories (SSP2 4.5, SSP3 7.0 and SSP5 8.5). Across all trajectories, model uncertainty peaks when the two ΔT_*K*_ values lie at the extremes of the considered range (1°C and 2.5°C), reinforcing the intuition that the more divergent the ecosystem responses, the less likely a fixed (no learning) strategy will perform well in both cases. Overall, model uncertainty remains lower under optimistic climate scenarios (averaging 1.2% for SSP1 1.9 and SSP1 2.6) than under pessimistic ones (2.5%, 2.1%, and 2.3% for SSP2 4.5, SSP3 7.0, and SSP5 8.5, respectively). This is due to the weaker non-stationarity of the most optimistic trajectories, reducing the importance of the ecosystem response to higher temperatures on decision-making.

**Figure 3:**
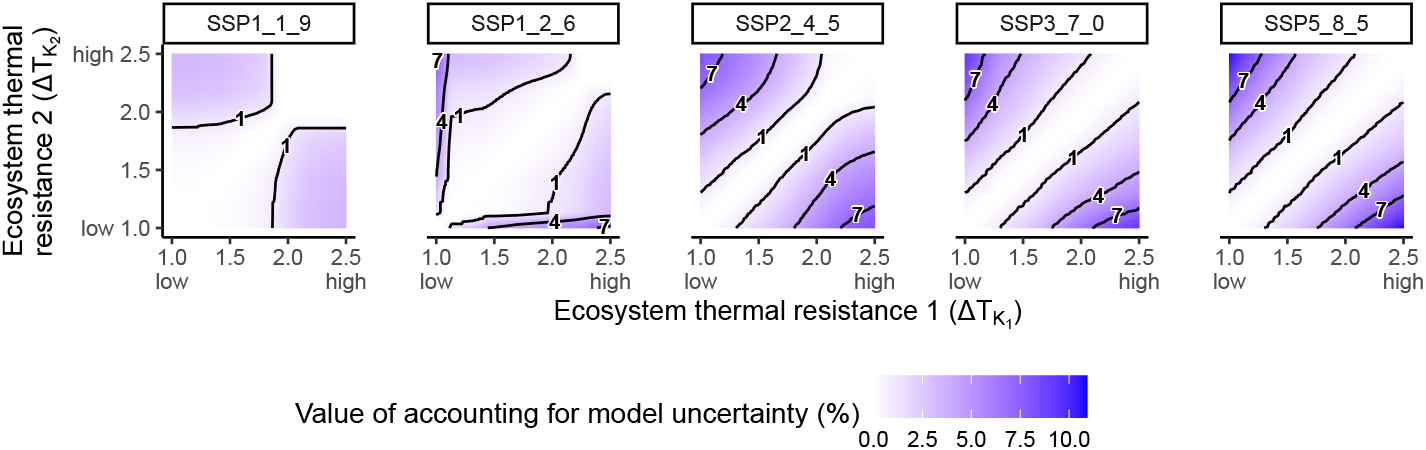
Value of accounting for model uncertainty, when the decision-maker is uncertain between two scenarios of thermal resistance (two competing values of ΔT_*K*_), across different climate change trajectories.

### 3.3 The value of modelling non-stationarity

For any non-stationary MDP, the value of non-stationarity depends primarily on the climate change trajectory. The more pessimistic the climate change scenario, the stronger the non-stationarity and therefore the higher the value of non-stationarity (Figure 4A). The value of non-stationarity is the lowest under the most optimistic climate change scenario (SSP1 1.9), with an average cost of stationarity of 0.3%. This result suggests that removing non-stationarity from the model reduces the expected benefits by only 0.3%. In contrast, as the climate trajectory becomes more pessimistic, the value of non-stationarity increases (15.4% on average under SSP5 8.5). These results reinforce the need to include non-stationarity for informed decision-making. The ecosystem’s response to climate change (ΔT_*K*_) also influences the value of non-stationarity (Figure 4B).

**Figure 4:**
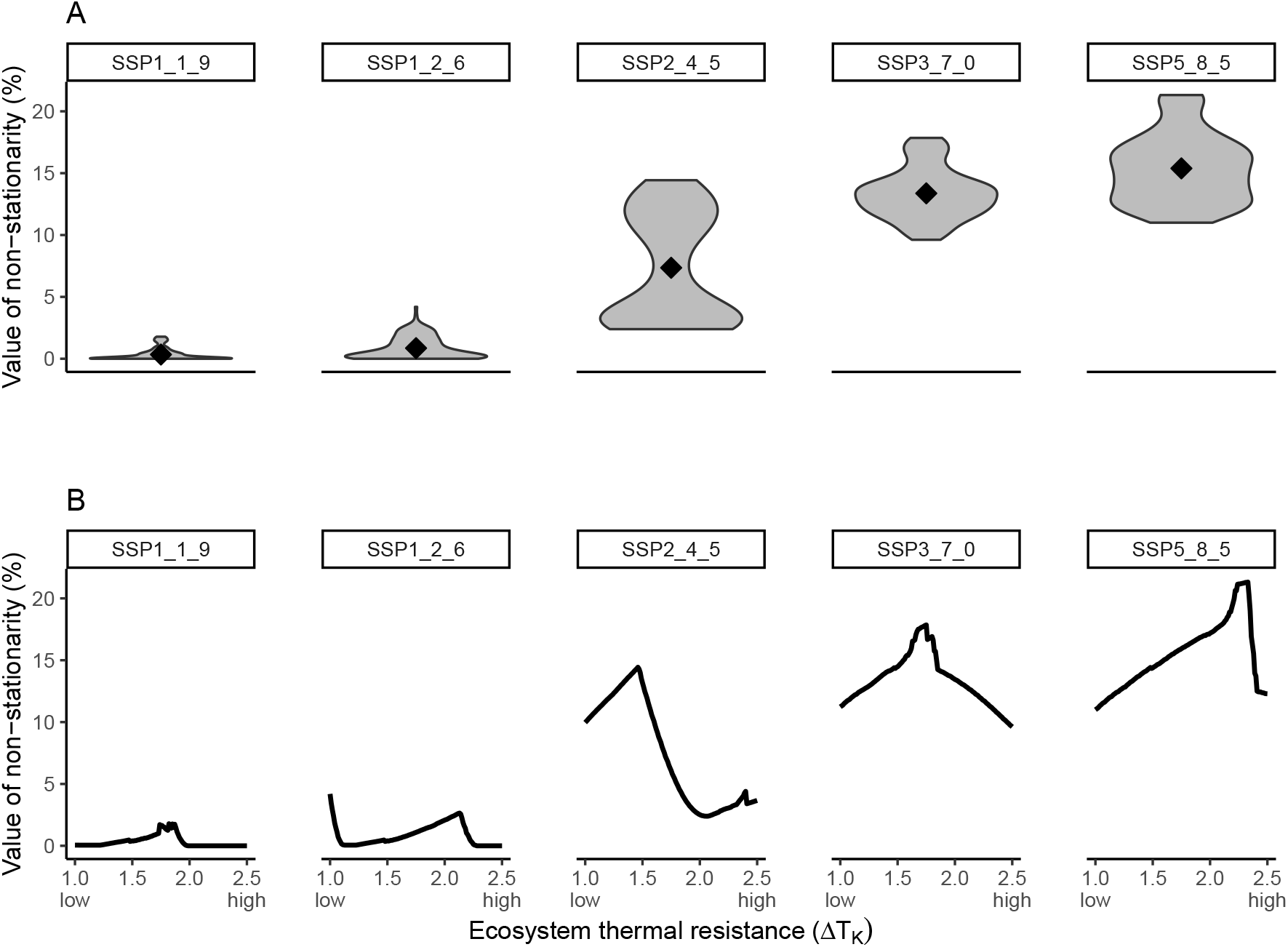
**A.** Summary of the value of modelling non-stationarity across different climate change scenarios. **B**. Value of modelling non-stationarity achieved for each possible scenario ecosystem response to climate change represented by the ecosystem thermal resistance (ΔT_*K*_).

### 3.4 Outputs of the protocol

We find that the possible modelling simplifications heavily depend on the climate change trajectory (Figure 5). For climate scenarios SSP1 1.9 and SSP1 2.6, the decision problem can be simplified to a stationary MDP (light blue area of Figure 5) if two conditions are met. First, both competing values of ΔT_*K*_ must fall within regions where the value of non-stationarity is low, for example, both values below 1.75°C or above 1.9°C for SSP1-1.9 (Figure 4B). Second, the two competing values must be sufficiently close to each other, such that the value of model uncertainty is also low (white region in Figure 3). If the second condition is not satisfied, the decision problem can often still be reduced to a stationary adaptive management problem (pink regions of Figure 5). Conversely, if non-stationarity is a valuable factor, but the competing trajectories remain close (near the 1:1 line in the dark blue region of Figure 5), the decision-problem can be simplified to a non-stationary MDP. In all other cases, where both the value of non-stationarity and of model uncertainty are high, our protocol recommends applying a non-stationary adaptive management approach.

**Figure 5:**
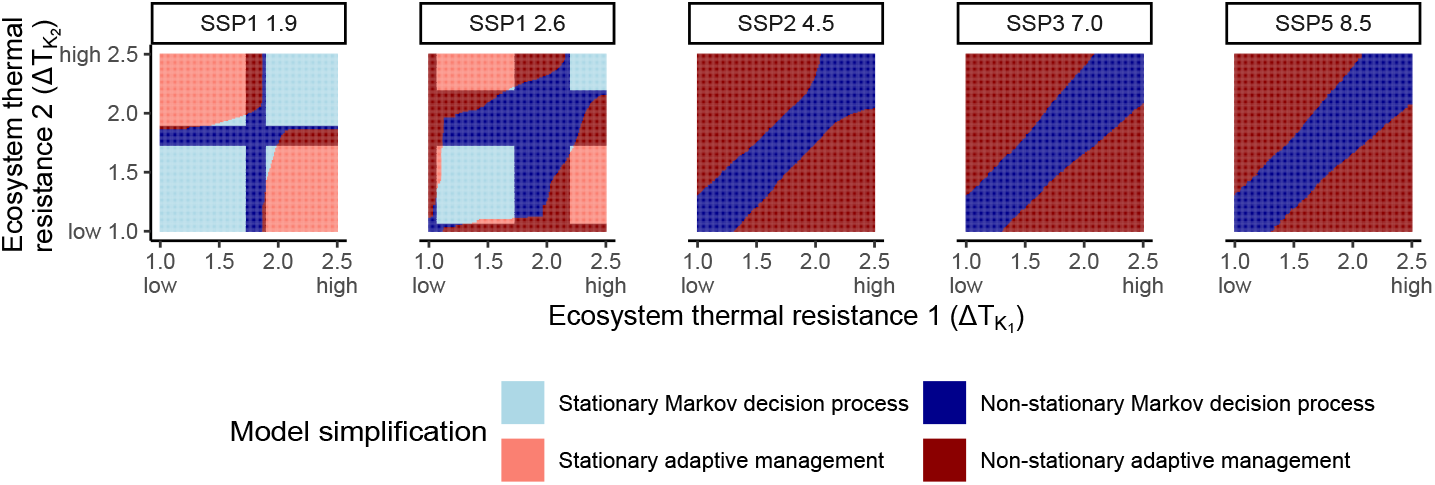
Model simplifications when the threshold in expected losses is 1% of total expected benefits. Here, the decision-maker has two competing scenarios for the ecosystem response to climate change, represented by the thermal resistance (ΔT_*K*_), and is uncertain about which one is true. For each combination of values and each climate change trajectory (assumed fully known), we summarize the model recommendation.

Interestingly, for the more pessimistic trajectories (SSP2 4.5, SSP3 7.0 and SSP5 8.5), the high value of non-stationarity consistently leads to the recommendation to explicitly include time as a state variable in the decision-making process. For these scenarios, when the competing values of ΔT_*K*_ are close enough, a non-stationary MDP still yields high benefits for the ecosystem (dark blue region of Figure 5 along the 1:1 line). However, when the difference between the two values of ΔT_*K*_ becomes too large, the expected losses become too important to justify simplifying the problem to a non-stationary MDP. In these situations, our protocol recommends a non-stationary adaptive management strategy (dark red region of Figure 5).

Logically, the size of each recommended model region varies depending on the acceptable loss threshold (Figure S2). A lower threshold leads to larger regions where the non-stationary adaptive management approach is recommended, while the regions where simpler decision approaches are sufficient become smaller (Figure S2A). Conversely, increasing the loss threshold reduces the area where non-stationary adaptive management is recommended, resulting in larger regions where simpler models can be applied. This makes sense because, by definition, a non-stationary adaptive management approach will always provide an optimal solution to this problem, while other approaches may not always do.

## 4 Discussion

We present a value of information protocol to guide decision-makers in managing ecosystems with uncertain and non-stationary dynamics. While the best management approach is non-stationary adaptive management (Keith et al., 2011; Chadès et al., 2017), solving such problems is computationally challenging (Chadès et al., 2012) and often produces solutions that are difficult to interpret and implement in practice (Dujardin et al., 2015, 2017; Ferrer-Mestres et al., 2021). To support decision-makers in simplifying their decision models, we use two metrics that quantify the value of modelling non-stationarity and uncertainty in ecosystem dynamics. This approach enables decision-makers to find a trade-off between the interpretability of simpler, suboptimal strategies and the performance of more complex and optimal methods. We demonstrate the application of this protocol through a case study on coral reef management using new technologies to mitigate climate change impacts, inspired by Australia’s Great Barrier Reef.

Our protocol identifies situations where the sequential decision problem can be simplified from a non-stationary adaptive management problem to a stationary MDP. Choosing a simpler model reduces computational complexity and leads to more interpretable recommendations for on-ground managers, which are more likely to be implemented (e.g. locally mitigating temperature by 1°C as long as coral cover remains positive; Figure 2). Our protocol also offers a quantitative justification to decide when to adopt a more complex, non-stationary adaptive management approach.

We find that the climate trajectory is the main driver determining whether the decision model can be simplified. As non-stationarity is highly valuable for decision-making for the more pessimistic climate trajectories (SSP2 4.5, SSP3 7.0 and SSP5 8.5), we found that the decision-model could at best be simplified to a non-stationary MDP, but generally requires a full non-stationary adaptive management approach. Conversely, as non-stationarity is less valuable in the most optimistic trajectories (SSP1 1.9 and SSP1 2.6), the decision model can often be reduced to a stationary MDP. These results remain consistent for different thresholds of acceptable losses (Figure S2).

Our value of information protocol using MDPs, can be extended to include multiple ecosystem health indicators (beyond coral cover) by extending the MDP state space. Our approach is also generalizable to address interacting, non-stationary factors such as water quality and seasonality, and uncertainty in the climate change trajectory. In this study, we modeled only one type of ecosystem response to climate change (a smooth decline) and explored problems with only two uncertain responses. This simplification allowed us to present a clear examples of how our protocol can be applied in practice. Future work could explore other responses, including abrupt, stepped or stochastic changes (Lindkvist et al., 2017; Bergstrom et al., 2021). In addition, our model implicitly assumes that ecosystems will always be able to recover with high temperature mitigation. In practice, ecosystems might hit degradation tipping points (Dakos et al., 2019), where restoration might no longer be possible, and causing the model to lose the Markov property. Future work could benefit from advances in AI to solve MDPs and POMDPs with large state and action space such as deep reinforcement learning (François-Lavet et al., 2018). Deep reinforcement learning can also accommodate to non-Markovian decision problems by inferring hidden past states, which allows to recover the Markov property (Lapeyrolerie et al., 2022). Finally, the decision-models recommended by our approach are MDPs or POMDPs, which solutions remain challenging to interpret in most cases. Future work could apply existing methods to improve the interpretability of MDP and POMDP solutions recommended by our approach (Ferrer-Mestres et al., 2021; Dujardin et al., 2015, 2017; Pascal et al., 2020).

This study contributes to the long-standing debate: when should adaptive management programs be used to manage dynamic, uncertain ecosystems? Our approach offers a transparent answer to this question, contributing to deepening our understanding on the value of information and adaptive management (Hauser and Possingham, 2008; Williams and Johnson, 2015; Tulloch et al., 2017; Holden et al., 2024) and extending it to non-stationary contexts. This contribution is relevant across many domains beyond ecology, including AI (Lecarpentier and Rachelson, 2019) and infrastructure adaptation (Bhattacharya et al., 2025; Feng et al., 2025), where an increasing number of studies incorporate non-stationary dynamics into decision-making, often to account for external factors such as climate change. Our method provides a clear framework for assessing whether accounting for non-stationarity is valuable for decision-making and demonstrates its potential to simplify complex decision-making processes.

Our findings have important implications for ecosystem management under climate change. By providing a structured, quantitative approach to model selection, our protocol supports decision-makers in choosing the simplest model that meets performance requirements, without compromising ecological outcomes. This is particularly valuable in practice, where optimal but complex models show a slower uptake due to the technical expertise required to find solutions and the difficulty in interpreting solutions. In addition, by explicitly identifying when non-stationarity or structural uncertainty cannot be ignored in decision-making, our approach prevents the risk of adopting misleadingly simple models, as we have shown for the more pessimistic climate trajectories. Approaches like ours are further needed to help bridge the gap between computer science and applied ecology.

## Acknowledgments

We thank Dr Ryan Heneghan and Nina Rynne for providing valuable insights into the different climate change scenarios. LVP and KJH were funded by Australian Research Council Discovery Early Career Researcher Award DE200101791. MPA’s contribution was funded by an Australian Research Council Discovery Early Career Resarcher Award DE200100683.

