## Supplemental Materials for "When should we use non-stationary adaptive management? A value of information analysis"

#### Supplementary material

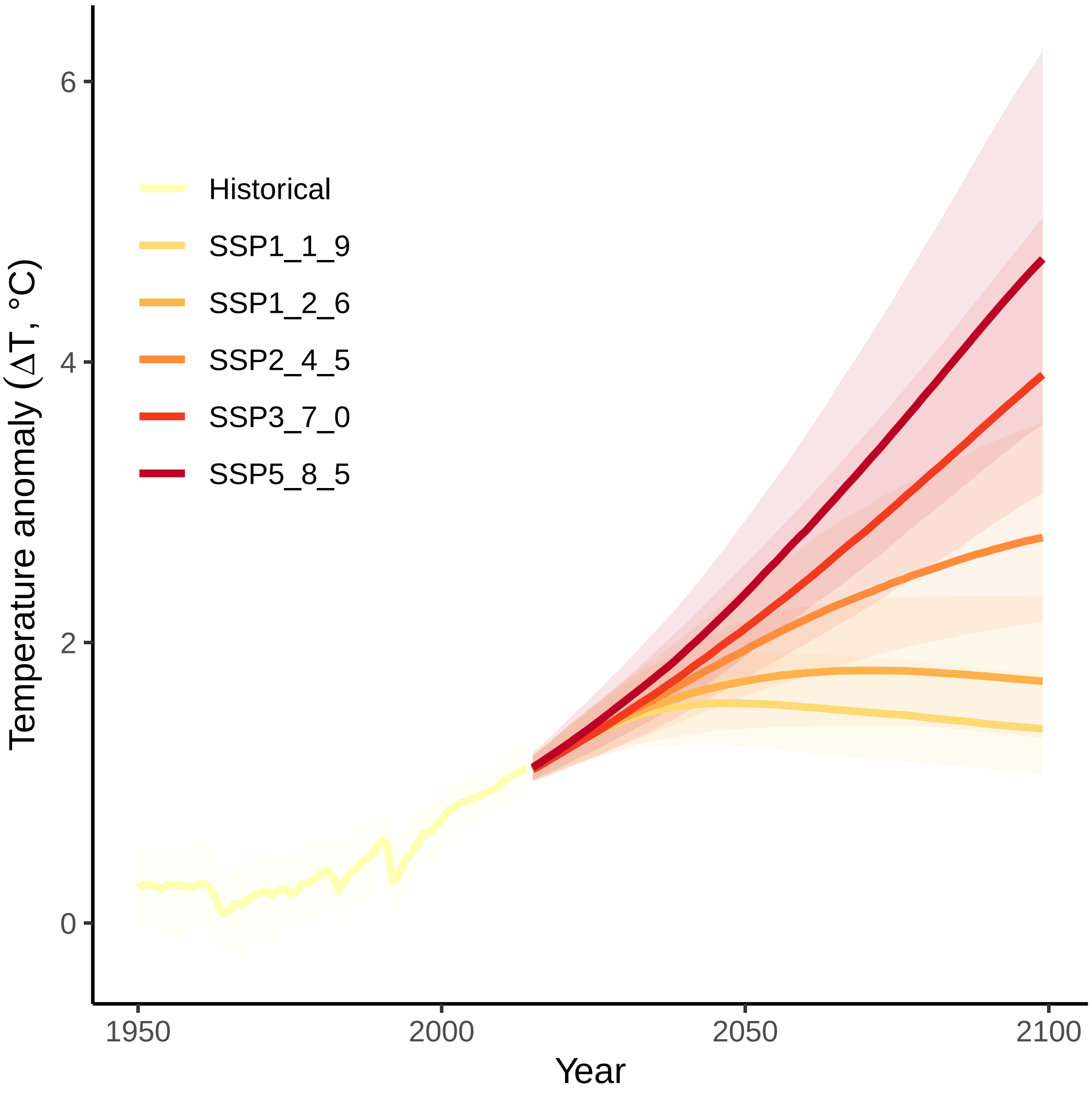


Figure S1: Historical and projected global mean surface air temperature anomalies relative to 1850-1900 (ΔT), according five climate change trajectories (IPCC, 2023).

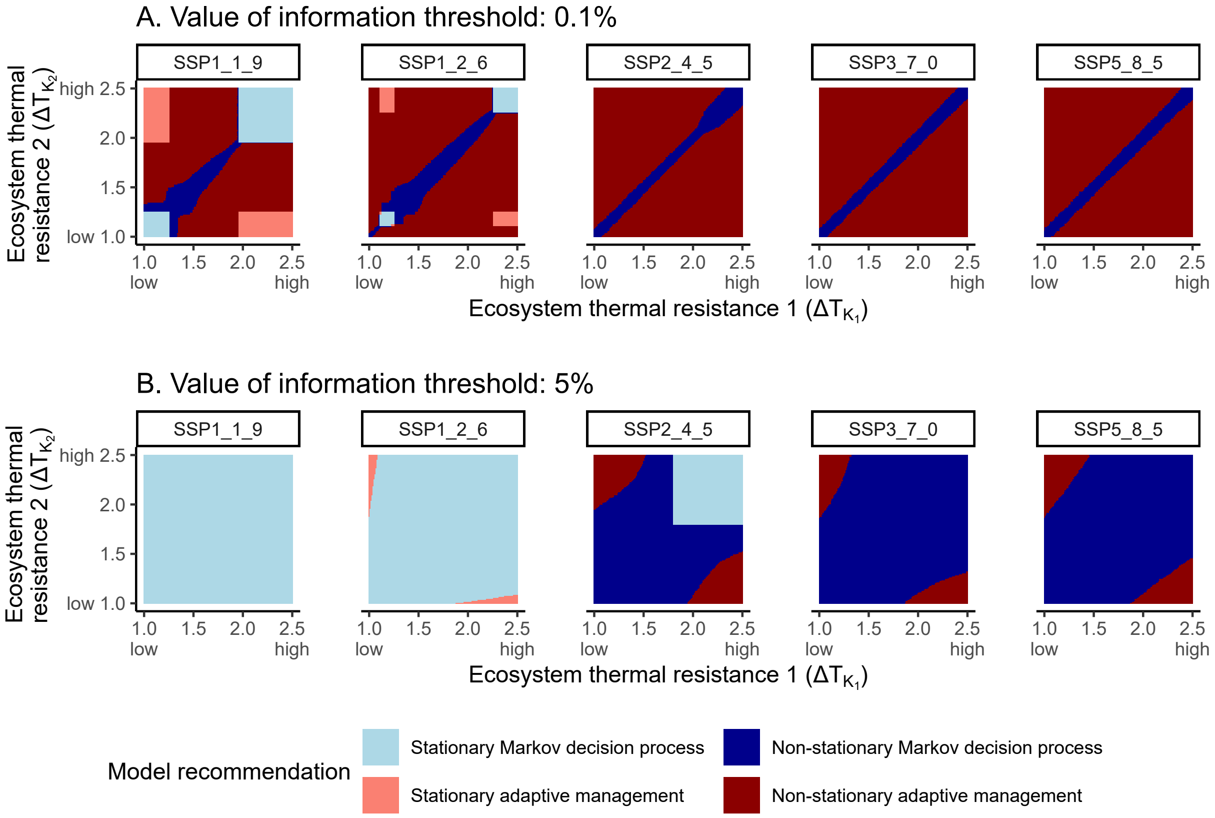


Figure S2: Model recommendations when the threshold in expected losses is 0.1% and 5% of total expected benefits. Here, the decision-maker has two competing scenarios for the ecosystem response to climate change, represented by the thermal resistance ($\Delta T_K$), and is uncertain about which one is true. For each combination of values and each climate change trajectory (assumed fully known), we summarize the model recommendation.

### A Non-stationary Markov Decision Processes

Markov Decision Processes (MDPs, Sigaud and Buffet (2013)) are a mathematical tool for sequential decision-making under uncertainty. Under complete knowledge of the ecosystem response to climate change ($\Delta T_K$), we find optimal recommendations for ecosystem management by framing the problem as a non-stationary MDP. For a given climate change trajectory, we denote $\Delta T_t$ the temperature anomaly at time $t$ (assumed fully known by the decision-maker). In this study, we use the average temperature anomaly for each trajectory, but recognize there is uncertainty in the projected temperatures as illustrated in Figure S1. For a given climate trajectory, the non-stationary MDP is then defined with:

- $X = S \times \Theta$ the factored set of states. $S$ represents the set of possible states of the ecosystem (coral cover), that we discretize into $N = 11$ states from 0 to 1 by 0.1 increments. $\Theta$ represents the set of time steps for decision-making. Using the projections for temperature anomalies (IPCC, 2023), we simulate decision-making between 2015 and 2100, therefore $\Theta = [2015, ..., 2100]$. The state of the system $x$ is fully described by an ecosystem state $s$ and a time step $t$;

- $A$ is the discrete set of possible actions, that correspond to the different pressure mitigation levels ($\Delta T^{tech}$). Here $A = [0, 1, 2]$. For clarity, we denote $\Delta T^{tech}$ the temperature mitigation level implemented;

- $P$ is the transition function between states at time $t$, where $P_t(s'|s, \Delta T^{tech})$ is the probability of transitioning from $s$ to $s'$ when the temperature mitigation is $\Delta T^{tech}$ and at time $t$. Since we have:

$$\bar{s_t} = s_t + s_t.r\left(1 - \frac{s_t}{K(\Delta T_t - \Delta T_t^{tech})}\right),$$
$$\frac{\log(s_{t+1}) - \log(\bar{s_t})}{\sigma} \sim \mathcal{N}(0, 1),$$

where:

$$K(\Delta T_t - \Delta T_t^{tech}) = K_{min} + (K_{max} - K_{min})\left(1 - \frac{1}{1 + e^{-m(\Delta T_t - \Delta T_t^{tech} - \Delta T_K)}}\right) \quad ,$$

then, the transition probability between states is :

$$P_t(s'|s, \Delta T^{tech}, y) = \int_{\max(0, s' - \frac{1}{2*N})}^{\min(1, s' + \frac{1}{2*N})} \mathrm{f}(x, \bar{s}_t, \sigma)\mathrm{d}x \quad .$$

where f is the density function of a log-normal distribution.

- $r_t$ is the reward function at time $t$, where $r_t(s, a)$ is the instant reward obtained at time $t$, when the system is in state $s$ and action $a$ is implemented. Here, we consider that the reward function is independent of time:

$$r(s, \Delta T^{tech}) = b.s - c.(\Delta T^{tech})^2$$

- $H$ is the horizon of the problem, here until 2100;

- $\gamma$ is the discount factor relating the relative importance of current rewards compared to future ones. We set $\gamma = 0.99$;

Solving a non-stationary MDP means finding the best sequence of actions to maximize the expected sum of discounted rewards. The sequence of actions is called a strategy, and is at each time step a strategy maps a state (combination of ecosystem state $s$ and time step $t$) to an action $\pi : S \times \Theta \to A$. We therefore aim to find:

$$V^*(s, t = 0) = \max_{\pi} \mathbb{E}\left(\sum_{i=0}^{H} \gamma^i r_i(s_i, \pi(s, i)) \middle| s_0 = s, t = 0\right). \quad \text{(S1)}$$

A non-stationary MDP can be solved using backward stochastic dynamic programming (Sigaud and Buffet, 2013):

$$\begin{cases} V^*(s, t) = \max_{a \in A} r_t(s, a) + \gamma \sum_{s' \in X} P_t(s'|s, a) V^*(s', t + 1), \\ V^*(s, H) = r_H(s, a). \end{cases} \quad \text{(S2)}$$

### B Non-stationary active adaptive management under model uncertainty

For each climate change scenario and each experiment, we define active adaptive management under model uncertainty as a Partially Observable Markov Decision Process with:

- $X = S \times \Theta$ the factored set of fully observable states, $A$ the discrete set of possible actions, $r$ the reward function, $H$ the horizon and $\gamma$ the discount factor are the same as in the non-stationary MDP (Section A);
- $Y$ is the finite set of non-observable states describing the possible non-stationary dynamics of the ecosystem. $Y$ is defined by the possible ecosystem responses to climate change, and is represented by different ecosystem resistance scenarios (different temperatures $\Delta\mathrm{T}_K$). For example in case study A, we consider two possible models: the first with $\Delta\mathrm{T}_K = 1$, and the second $\Delta\mathrm{T}_K = 1.6$. Because the elements of $Y$ cannot be observed, we have a belief $b_t(y)$ over each model $y$. $b_t(y)$ is the probability that the model $y$ is the true model at time $t$;
- $P_t$ the transition function between states at time $t$, where $P_t(s'|s, \Delta\mathrm{T}^{\mathrm{tech}}, y)$ is the probability of transitioning from $s$ to $s'$ when the temperature is mitigated by $\Delta\mathrm{T}^{\mathrm{tech}}$ and at time $t$ and according to model $y$. $P_t$ has the same definition as for non-stationary MDP with a different values of $\Delta\mathrm{T}_K$ for each model;
- $P_t^y$ represents the transition function between models at time $t$. Here we assume that the system cannot transition from one model to the other, therefore $P_t^y$ is the identity matrix;
- the set of observations are the observable states: the time step and the ecological state, and the observation function is the identity matrix;

Because the set of possible models $Y$ is not observable, we need to infer the true model at each time step based on the ecosystem response to climate change. We use Bayes' rule for POMDPs to update the belief as follows (Chadès et al., 2012). At time $t$, the decision-maker has a belief $b_t$ over each possible model and the ecosystem state is $s$. If the manager implements the action $\Delta\mathrm{T}^{\mathrm{tech}}$, and observes that the ecosystem state changes to $s'$, then the belief is updated to $b_{t+1}$ as:

$$b_{t+1}(y) = \frac{P_t(s'|s, \Delta\mathrm{T}_t^{\mathrm{tech}}, y) b_t(y)}{\sum_{\tilde{y}} P_t(s'|s, \Delta\mathrm{T}_t^{\mathrm{tech}}, \tilde{y}) b_t(\tilde{y})} \quad . \tag{S3}$$

At each time step, the manager must decide what action to execute given the current state of the ecosystem $s_t$ and current belief $b_t$. Solving a POMDP means finding a strategy $\pi$ that maximizes a criterion: the expected discounted sum of long-term benefits. A strategy $\pi$ is defined as a function mapping an ecosystem state, a time step and a belief state to an action (or $\pi : s \times \Theta \times B \to A$). POMDP solvers aim to find a strategy such that:

$$V^*(s,t,b) = \max_{\pi} \mathbb{E}\left( \sum_{i=0}^{H} \gamma^i r(s_i, \pi(s,i,b_i)) \middle| s_0 = s, t = 0, b_0 = b \right) \quad ,$$

where $V^*$ is called the optimal value function of a this POMDP.
